# Characterization of Stress Tolerance and Probiotic Traits of *Streptococcus salivarius* Strains Isolated from Wildflower Honey

**DOI:** 10.64898/2026.09.27.754824

**Authors:** Wonhui Lee, Yeon-Soo Oh, Chan-Soo Ock, Ye-Eun Oh, Ha-young Kim, Hyung-Ki Do, Chul-Won Hwang

**Author notes:** Corresponding authors: Hyung-Ki Do and Chul-Won Hwang.

## Abstract

Honey represents a distinctive microbial habitat that imposes physicochemical stresses on its resident microbiota while also serving as a potential source of beneficial microorganisms. Two *Streptococcus salivarius* strains, DOW-5A and DOW-10, isolated from Korean wildflower honey were identified through phenotypic observation and 16S rRNA sequencing. Comprehensive evaluation, including 5 safety assessments and 14 functional assays, was conducted to reveal distinct potential probiotic characteristics in the two isolates. Both strains exhibited no gelatinase or hemolytic activity and showed no significant cytotoxicity toward Caco-2 cells. While DOW-5A maintained higher viable counts during the simulated gastric phase, DOW-10 showed an increase after an initial decline during the simulated intestinal phase. Both strains also exhibited growth under stress conditions involving phenol or hydrogen peroxide. Among the cell-surface and adhesion-related characteristics, DOW-5A exhibited greater mucin adhesion and cell-surface hydrophobicity, whereas DOW-10 showed higher autoaggregation at 6 and 12 h. Both strains exhibited DPPH radical-scavenging activity, with no significant difference from that of the reference probiotic strain, *Lactobacillus rhamnosus* GG. Atypical and crystalline-like colony morphologies, suggestive of EPS production, were observed when sucrose was supplied as the carbon source. Furthermore, both strains formed biofilms in a crystal violet staining assay, while DOW-5A showed a marked increase in biofilm biomass upon sucrose supplementation. These findings highlight strain-specific variation in stress tolerance and probiotic-associated traits between the two honey-derived *Strep. salivarius* isolates and support their further investigation as probiotic candidates.

## 1. Introduction

Honey is a natural product well known for its health-promoting activities, including antimicrobial, antioxidant, and anti-inflammatory activities [1]. Its more than 200 components, including antimicrobial substances such as phenolic compounds, hydrogen peroxide (H₂O₂), and organic acids [2, 3], contribute to its inhibitory effects on microorganisms alongside its physicochemical characteristics. Honey’s high osmotic pressure, low pH, and low water activity create extreme conditions for microbial growth [4, 5], making the internal environment harsh for microorganisms [6]. Indigenous microbial communities and the microbes introduced into honey must tolerate these stressful conditions. Several studies have suggested that these environmental constraints limit those that cannot adapt to this unique environment, thereby selecting for microorganisms resistant to various stresses [7, 8]. The microbial community of honey is particularly associated with the honeybee gut microbiota and nectar-producing plants [9]. The honey microbiota, shaped by these ecological processes, is a complex component of honey that may substantially influence its functional properties [10].

Among them, lactic acid bacteria (LAB) have been widely studied for their tolerance to various environmental stresses, and metagenomic analyses have confirmed their presence in honey flowers, the honeybee gut, and honey itself [11–13]. Recent studies have shown that some honey-derived LAB strains have high potential to withstand stressful conditions through metabolites such as exopolysaccharides (EPS) and to proliferate in sugar-rich environments [14, 15]. These characteristics may be relevant to probiotic potential, particularly the ability to survive stressful conditions encountered during gastrointestinal (GI) transit [16]. Even though traditionally used probiotics mainly belong to the genera *Lactobacillus* and *Bifidobacterium* [17], increasing attention has recently been given to *Streptococcus salivarius*, a Gram-positive LAB that colonizes the human oral cavity early in life [18, 19]. Several strains, including *Strep. salivarius* K12, have been investigated for health-promoting properties relevant to the oral cavity and GI environment [20–22]. Even though studies of the probiotic potential of *Strep. salivarius* have primarily focused on human-derived strains [23], there were several cases reported from non-human sources, including marine fish, fermented foods, terrestrial livestock, olive drupes, and even in the arctic driftwood [24–28]. These previous studies indicate the ecological distribution of *Strep. salivarius* is not restricted to human-associated environments. The objective of this study was to isolate LAB strains from Korean wildflower honey samples and to evaluate the phenotypic responses to physiological stresses and potential probiotic properties of two isolates identified as *Strep. salivarius*.

## 2. Materials and Methods

### 2.1. Isolation and Identification

#### 2.1.1 Sample Preparation and Isolation of LAB

In this study, wildflower honey samples collected from Jirisan National Park, located in Sancheong, South Korea were used and stored at room temperature. All procedures were strictly carried out under aseptic conditions. The samples were serially diluted with sterile phosphate-buffered saline (PBS), then spread onto De Man, Rogosa, and Sharpe (MRS) agar supplemented with 0.5% (w/v) CaCO₃ and incubated at 37°C for 24 h [29]. The colonies presumed to be LAB were selected based on clear zone formation, colony morphology, Gram staining, and catalase activity. These isolates were repeatedly streaked onto MRS agar, and the pure cultures were mixed in equal volumes with a 50% (v/v) glycerol solution and stored at −80°C. The reference strain, *Lacticaseibacillus rhamnosus* GG (ATCC 53103, LGG), was provided by Dr. Bobae Kim (Handong Global University) and used as a reference strain [30].

#### 2.1.2 Molecular Identification of LAB Isolates

To characterize the isolates at the molecular level, 16S rRNA gene amplification and sequencing were performed by Macrogen Inc. (Seoul, South Korea). The obtained sequences were compared with reference sequences in GenBank using the Basic Local Alignment Search Tool at the National Center for Biotechnology Information (NCBI). The 16S rRNA gene sequences of strains DOW-5A and DOW-10 were deposited in GenBank.

#### 2.1.3 Phylogenetic Analysis

Sequences of the isolates and reference strains were aligned using MEGA version 12 [31]. Using the maximum-likelihood method with the Kimura two-parameter model, the phylogenetic tree was constructed, and the reliability of the tree was assessed through 1,000 bootstrap replicates.

### 2.2 Safety Assessment

#### 2.2.1. Gelatinase and Hemolytic Activities

The isolates were inoculated into nutrient gelatin medium (MBcell, Seoul, Korea) and incubated at 37°C for 24 h. The media were then cooled at 4°C for 1 h and were observed for liquefaction. Brain Heart Infusion agar (Difco, Sparks, MD, USA), supplemented with 5% (v/v) defibrinated sheep blood, was used to evaluate the hemolytic activity. After incubation at 37 °C for 24 h, hemolysis was classified as α-hemolysis (partial hemolysis), β-hemolysis (complete hemolysis with a clear zone), or γ-hemolysis (no hemolysis). *Staphylococcus aureus* (KCTC 1916) was used as the positive control in both assays [32].

#### 2.2.2. Antibiotic Susceptibility Testing

LSM, consisting of 90% (v/v) Iso-Sensitest broth (Oxoid, UK) and 10% (v/v) MRS broth, was used as the test medium. The types of antibiotics were ampicillin (0.032–16 μg/mL), vancomycin (0.25–128 μg/mL), gentamicin (0.5–256 μg/mL), streptomycin (0.5–256 μg/mL), erythromycin (0.016–8 μg/mL), clindamycin (0.032–16 μg/mL), and tetracycline (0.125–64 μg/mL). Strains cultured on MRS agar at 37°C for 24 h were suspended in LSM to a concentration of 1 × 10⁶ CFU/mL, then mixed 1:1 (v/v) with LSM containing a two-fold concentration. The wells without added antibiotics were used as the growth control. The MIC was defined as the lowest antibiotic concentration that completely inhibited visible growth after 24 h of incubation at 37°C [33].

#### 2.2.3. Caco-2 Cell Culture

Human colorectal adenocarcinoma Caco-2 cells (HTB-37) were provided by Dr. Jung-Min Lee at Handong Global University. The cell line was not independently authenticated prior to this study. The cells were maintained in RPMI-1640 medium (Gibco, Grand Island, NY, USA) supplemented with 10% (v/v) fetal bovine serum (FBS; Gibco) and 1% (v/v) penicillin–streptomycin (Gibco) at 37 °C in a humidified atmosphere containing 5% CO₂. The culture medium was replaced every 2–3 days.

#### 2.2.4. Live/Dead Cell Staining Assay

Cell viability and membrane integrity were assessed using a LIVE/DEAD Viability/Cytotoxicity Kit (Invitrogen, Carlsbad, CA, USA) [34]. Following cell-free supernatant (CFS) treatment for 24 or 48 h, cells were washed twice with Dulbecco’s phosphate-buffered saline (DPBS) and stained with 2 μM calcein-AM (green fluorescence, viable cells) and 4 μM ethidium homodimer-1 (EthD-1; red fluorescence, dead cells) for 30 min at room temperature in the dark. Fluorescence images were acquired using a fluorescence microscope (Leica Microsystems, Wetzlar, Germany).

#### 2.2.5. Cell Viability Assay (CCK-8)

The CFS of two strains was used to evaluate the cytotoxicity in Caco-2 cells using a Cell Counting Kit-8 (CCK-8; Dojindo Laboratories, Kumamoto, Japan). Cells were seeded in 96-well plates at 1 × 10⁴ cells/well and incubated for 24 h, followed by treatment with various concentrations of CFS for 24 or 48 h, with untreated cells serving as the control group. After treatment, cells were washed with DPBS and incubated with 100 μL of RPMI-1640 containing 10% (v/v) CCK-8 reagent for 2 h at 37°C in the dark. OD_450_ was measured, and cell viability was calculated as follows [34]:

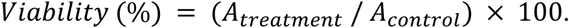

A_treatment_ and A_control_ represent the OD_450_ values of the sample and control.

### 2.3. GI Survival and Stress Tolerance

#### 2.3.1. Survival under Simulated GI Conditions

Survival under simulated GI condition was performed with reference to the INFOGEST 2.0 protocol [35]. The strains were cultured at 37°C for 24 h and then centrifuged at 4°C and 7,600 × g for 10 min. The recovered cells were washed with PBS and adjusted to a concentration of 1.0 × 10⁷ CFU/mL. After each stage, the samples were centrifuged at 4°C and 8,000 × g for 2 min, washed twice with PBS and resuspended in the solution for the next stage. For the oral phase, simulated salivary fluid (SSF) was supplemented with lysozyme (2,000 U/mL) and CaCl₂ (1.5 mM) and adjusted to pH 7.0. The samples were then incubated in SSF at 37 °C and 150 rpm for 2 min. For the gastric phase, simulated gastric fluid (SGF) was supplemented with pepsin (2,000 U/mL) and CaCl₂ (0.15 mM) and adjusted to pH 3.0. The samples were incubated in SGF at 37 °C and 75 rpm for 2 h. For the intestinal phase, simulated intestinal fluid (SIF) was supplemented with 0.3% (w/v) Bacto Oxgall (Difco, USA) and CaCl₂ (0.6 mM) and adjusted to pH 7.0. The samples were incubated in SIF at 37 °C and 75 rpm for 2 h. The samples were collected at the end of each stage to measure the viable cell count (log CFU/mL) and the absorbance at 600 nm.

#### 2.3.2. Acid and Bile Tolerance

MRS broth adjusted to pH 2.5 and MRS broth containing 0.3% (w/v) oxgall were prepared. Cultures grown at 37 °C for 24 h were inoculated into each medium at 1% (v/v) and incubated at 37 °C for 4 h. OD₆₀₀ was measured at 1, 2, 3, and 4 h, with untreated MRS broth used as the control. Relative growth was calculated at each time point as follows [36]:

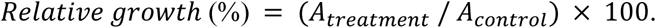

A_treatment_ and A_control_ represent the OD₆₀₀ values of the sample and control.

#### 2.3.3. Phenol and Hydrogen Peroxide (H₂O₂) Tolerance

Cultures grown in MRS broth at 37 °C for 24 h were inoculated at 1% (v/v) into MRS broth containing phenol (0.1, 0.2%, v/v) or H₂O₂ (1.0, 2.0 mM). OD₆₀₀ was measured at 6, 12, and 24 h, with untreated MRS broth used as the control. Relative growth was calculated at each time point as follows [37]:

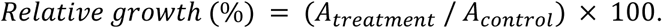

A_treatment_ and A_control_ represent the OD₆₀₀ values of the sample and control.

### 2.4. Adhesion-Related Properties

#### 2.4.1. Mucin Adhesion

Porcine mucin (0.5 and 1.0%, w/v) dissolved in PBS was added to 96-well plates (100 μL/well) and incubated at 4 °C for 12 h. The wells were washed twice with PBS. Cultures grown at 37 °C for 24 h were harvested by centrifugation at 8,000 × g for 2 min at 4 °C, washed three times with PBS, and adjusted to 1.0 × 10⁷ CFU/mL. 100 μL of cell suspensions were added to the mucin-coated wells and incubated at 37 °C for 2 h. The wells were then washed three times with PBS, and adherent cells were released with 100 μL of 0.1% (v/v) Triton X-100. The recovered cells were plated onto MRS agar and incubated at 37°C for 24 h before measuring the viable cell count (CFU/mL). The mucin adhesion rate was calculated as follows [38]:

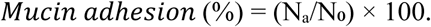

N₀ and Nₐ represent the initial number of viable cells and the number of viable cells recovered after washing.

#### 2.4.2. Autoaggregation

Cultures grown in MRS broth at 37 °C for 24 h were centrifuged at 8,000 × g for 2 min at 4 °C. The cell pellets were washed three times with PBS and adjusted to 1.0 × 10^7^ CFU/mL. 2 mL of cell suspensions were incubated without agitation at 37°C, and the OD₆₀₀ of the supernatant was measured at 3, 6, and 12 h. The auto-aggregation rate was calculated as follows [39]:

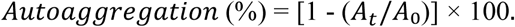

A₀ is the initial OD_600_ and Aₜ is the OD₆₀₀ of the upper suspension at each time point.

#### 2.4.3. Cell Surface Hydrophobicity

Cultures grown at 37 °C for 24 h were centrifuged at 8,000 × g, 4 °C for 2 min. The pellets were washed three times with PBS and adjusted to 1.0 × 10^7^ CFU/mL. 3 mL of cultures were separately mixed with 1 mL of the toluene (nonpolar solvent), chloroform (acidic solvent), and ethyl acetate (basic solvent). The mixtures were left 30 min for phase separation, and the OD_600_ of the aqueous phase was measured. The hydrophobicity was calculated as follows [40]:

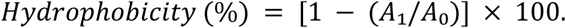

A₀ and A₁ represent the OD_600_ values before the solvent addition and after phase separation.

### 2.5. Functional evaluation

#### 2.5.1. Exopolysaccharide (EPS) Production

Cultures grown at 37 °C for 24 h were inoculated into MRS agar and MRS broth supplemented with 5% (w/v) sucrose. The inoculated media were incubated at 37°C for 24 h. After incubation, the cultures were visually examined for EPS-related phenotypes [41].

#### 2.5.2. 2,2-Diphenyl-1-picrylhydrazyl (DPPH) Radical-Scavenging Activity

All experimental procedures were conducted in the dark. Cultures grown at 37 °C for 24 h were adjusted to 1.0 × 10⁷ CFU/mL. 800 μL of culture supernatant was mixed with 1 mL of the 0.2 mM DPPH in methanol and reacted for 30 min at room temperature. The samples were centrifuged at 4°C and 12,000 rpm for 5 min, and 200 μL of the supernatant was dispensed into a 96-well plate to measure the OD_517_. 0.2 mM ascorbic acid solution prepared with DW was used as positive control. DPPH radical-scavenging activity was calculated as follows [42]:

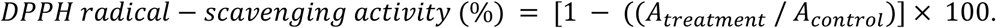

A_treatment_ and A_control_ represent the OD_517_ values of the sample and untreated control.

#### 2.5.3. Biofilm Formation

Cultures were grown in MRS broth at 37°C for 24 h and adjusted to 1.0 × 10⁷ CFU/mL, and 20 μL of the cultures were inoculated to 2 mL of MRS broth with and without 1% sucrose in the 24-well plates and incubated at 37 °C for 24 h. Uninoculated medium served as the blank. After removal, the wells were washed three times with PBS, air-dried, stained with 1% crystal violet (CV) for 20 min, and washed again. Biofilm-bound CV was solubilized with 33% (v/v) acetic acid, diluted 10-fold, and measured at 570 nm. Corrected OD₅₇₀ values were obtained by subtracting the corresponding medium blank. The relative change in biofilm formation resulting from the addition of sucrose was calculated as follows [43]:

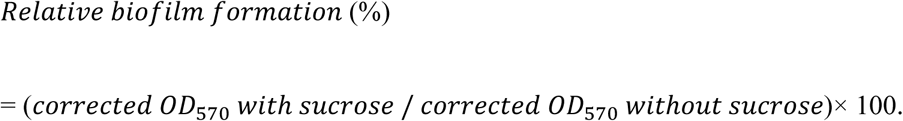

#### 2.5.4. Amylase and Protease Activities

Cultures were grown in MRS broth at 37°C for 24 h and adjusted to 1.0 × 10⁷ CFU/mL. 70 μL of the suspension was applied to 8 mm paper discs placed on International Streptomyces Project medium 4 agar (ISP 4; MBcell, Korea) for amylase or skim milk agar containing 1.5% (w/v) skim milk for protease. After incubation at 37 °C for 24 h, amylase activity was detected by iodine staining, and protease activity was determined by clear-zone formation around the discs [44].

### 2.6. Statistical Analysis

Statistical analyses were performed using GraphPad Prism version 8.0.2 (GraphPad Software, San Diego, CA, USA). All experiments were performed independently three times (n = 3), and results are expressed as mean ± standard deviation (SD). Differences among three or more groups involving a single factor were analyzed using one-way ANOVA. Two-way ANOVA was used for data involving two factors, and repeated-measures two-way ANOVA was used when the same experimental units were measured repeatedly over time. All ANOVA analyses were followed by Tukey’s multiple comparisons test. Direct comparisons between two strains were performed using an unpaired two-tailed Student’s t-test. Statistical significance was set at P < 0.05. Results of Tukey’s test are indicated by lowercase letters (a, b, c), assigned in descending order of group means within each comparison set. Groups sharing at least one letter did not differ significantly.

### 2.7. Nucleotide Sequence Accession Numbers

GenBank accession numbers PZ626003 and PZ626004 were assigned to the 16S rRNA sequences of DOW-5A and DOW-10, and will be publicly available upon publication.

## 3. Results

### 3.1. Isolation and Molecular Identification of LAB Strains

The overall workflow of this study is shown in Fig. 1. The isolates were Gram-positive and catalase negative (Supplementary Figure S1). Subsequently, the 16S rRNA sequences obtained from the two isolates were compared with four reference sequences available in GenBank. Both strains showed 99% sequence identity to *Strep. salivarius*. Based on these sequences, a phylogenetic tree was constructed, in which DOW-5A and DOW-10 clustered within the same clade as the reference strain *Strep. salivarius* ATCC 7073T (Fig. 2).

**Fig. 1.**
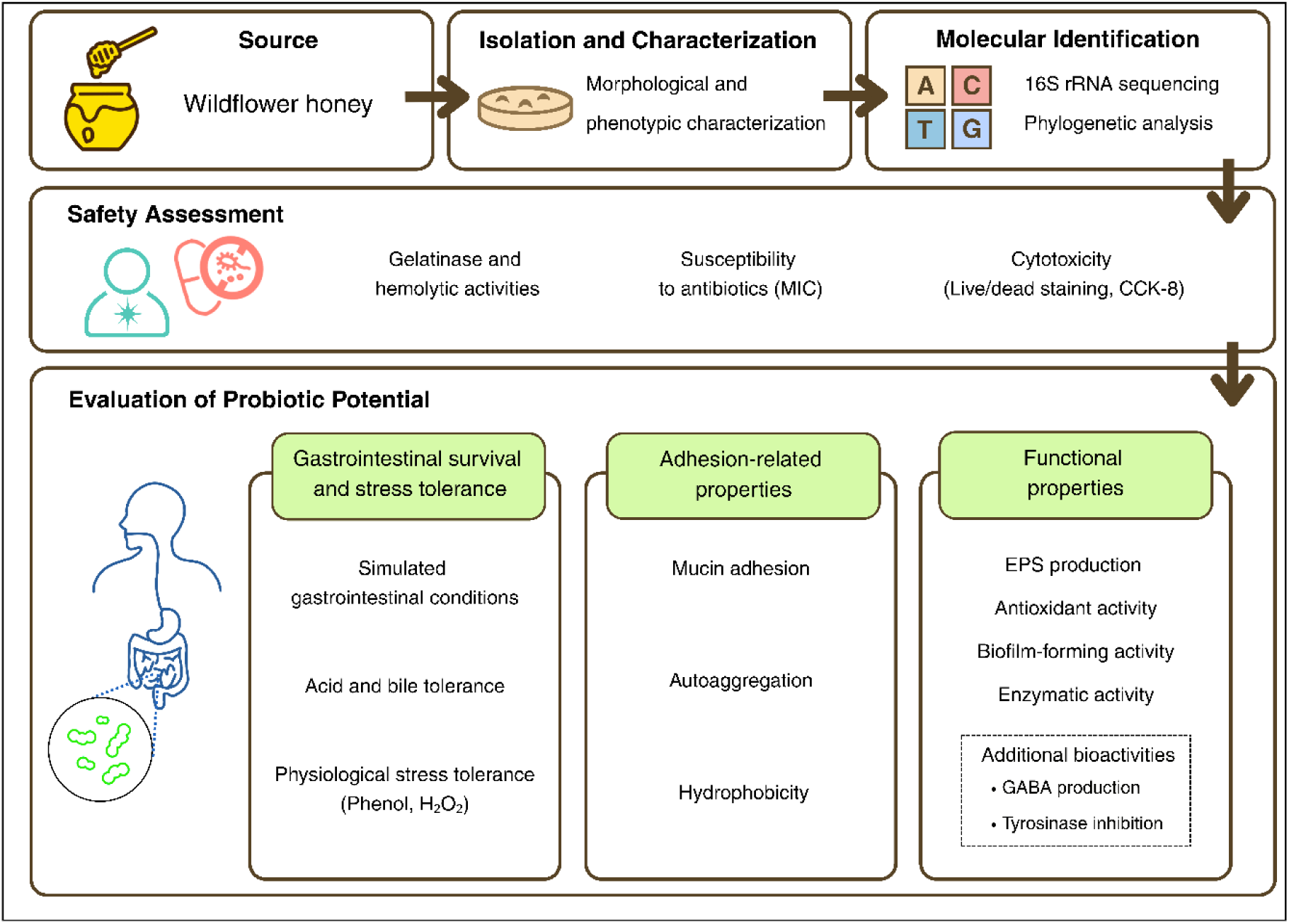
Schematic overview of the experimental workflow used for isolation, identification, and characterization of potential probiotic LAB from wildflower honey collected in Jirisan National Park, South Korea

**Fig. 2.**
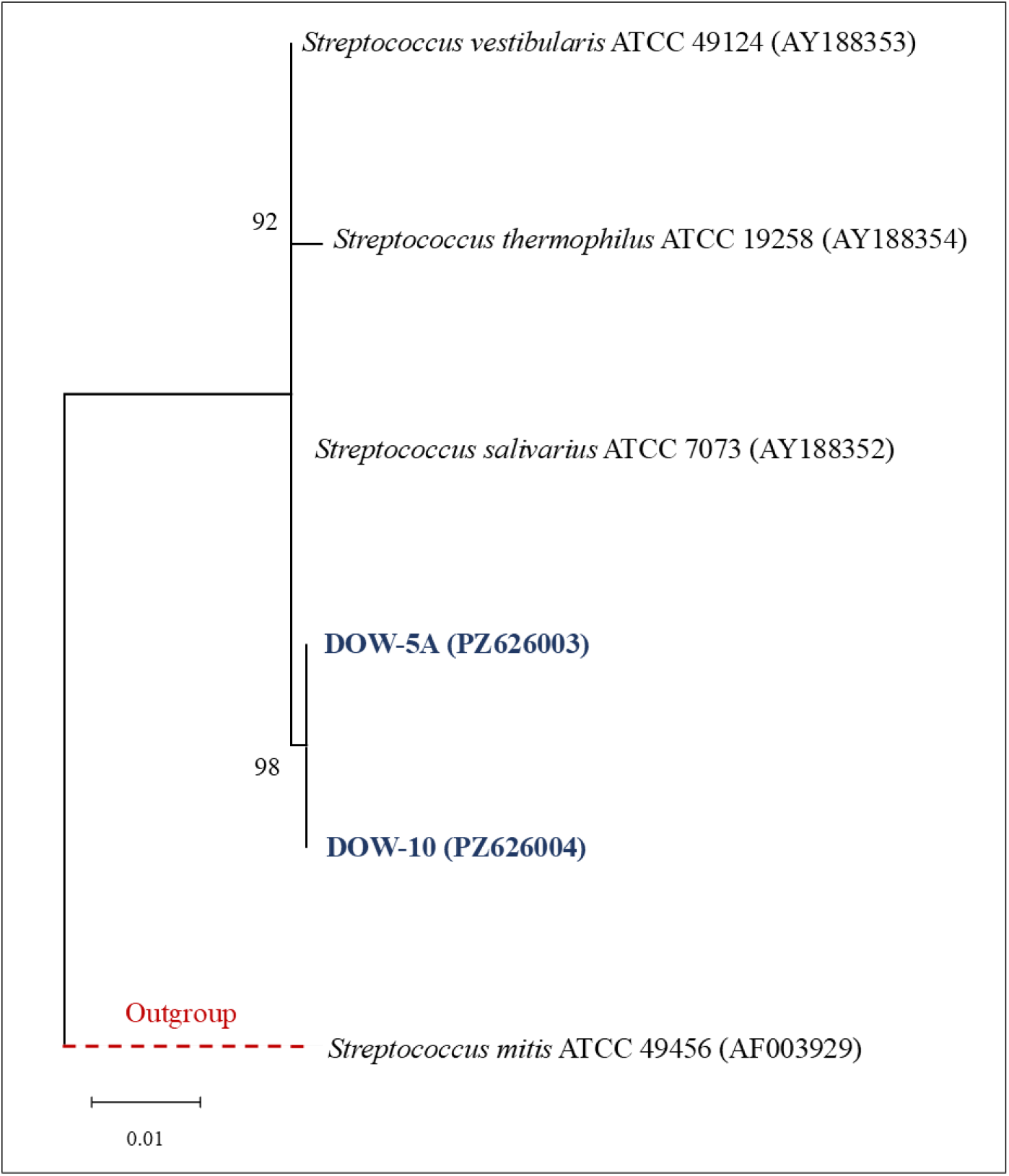
Phylogenetic tree of DOW-5A and DOW-10 based on aligned 16S rRNA sequences. Bootstrap values are indicated at the nodes

### 3.2. Safety Assessment

#### 3.2.1. Gelatinase and Hemolytic Activity

Both strains were negative for gelatinase activity. Neither strain produced a zone of hemolysis on BHI agar supplemented with 5% defibrinated sheep blood, classified as γ-hemolytic.

#### 3.2.2. Antibiotic Susceptibility (MIC)

The two strains showed different MIC profiles for the tested antibiotics (Table 1). DOW-5A and DOW-10 exhibited identical MIC values for ampicillin, gentamicin, streptomycin, clindamycin, and tetracycline, whereas their MIC values differed for vancomycin and erythromycin. In particular, DOW-10 showed a lower MIC for erythromycin than DOW-5A.

**Table 1.** MICs of seven antibiotics of DOW-5A and DOW-10.

| Strain | MIC (µg/mL) |  |  |  |  |  |  |
| --- | --- | --- | --- | --- | --- | --- | --- |
|  | AMP | VAN | GEN | STR | ERY | CLI | TET |
| DOW-5A | 4 | 4 | 32 | 64 | 2 | 0.5 | 0.5 |
| DOW-10 | 4 | 2 | 32 | 64 | 0.032 | 0.5 | 0.5 |

The experiments were independently performed three times, with each sample measured in triplicate in each experiment (n=3). AMP: ampicillin, VAN: vancomycin, GEN: gentamicin, STR: streptomycin, ERY: erythromycin, CLI: clindamycin, TET: tetracycline.

#### 3.2.3. Cytotoxicity in Caco-2 Cells

Both strains showed higher Caco-2 cell viability than positive control. Live/dead staining showed that most Caco-2 cells remained viable after 24 h of treatment with the CFS of both strains. In contrast, dead-cell staining was observed prominently in the positive control (Fig. 3a). Consistent with these findings, CCK-8 assay showed that after 48 h of treatment DOW-5A (88.21 ± 1.47%) and DOW-10 (89.88 ± 0.96%) maintained significantly higher cell viability than the positive control (20.82 ± 1.56%; P < 0.0001) (Fig. 3b).

**Fig. 3.**
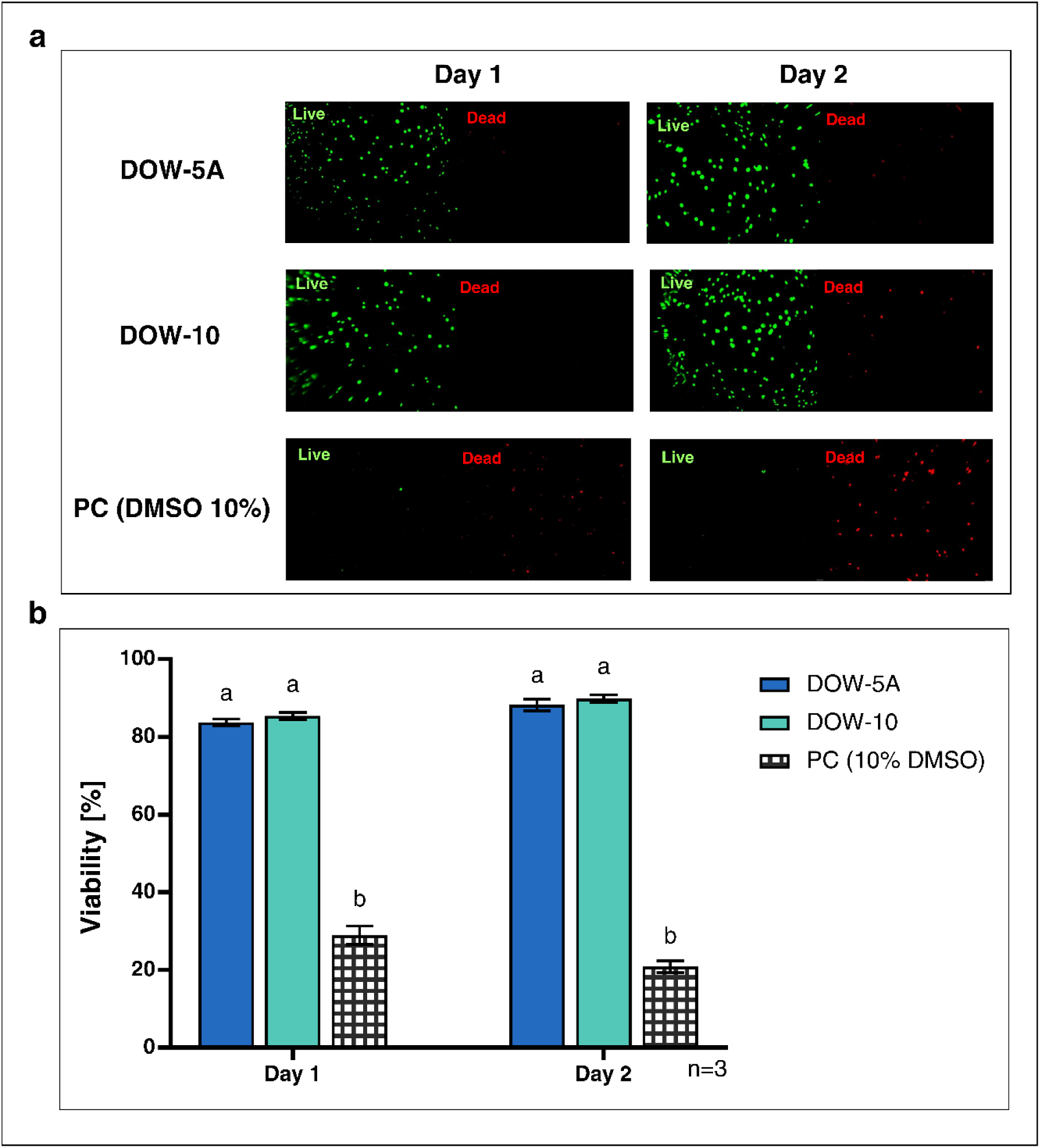
Cytotoxicity assays for the two strains were performed using (a) the live/dead assay and (b) the CCK-8 assay. All experiments were conducted over DAY 1 (24 h) and DAY 2 (48 h), and 10% DMSO was used as positive control. Three independent experiments were performed (n=3). Data were assessed by repeated-measures two-way ANOVA followed by Tukey’s multiple-comparisons test. PC: Positive control

### 3.3. GI Survival and Stress Tolerance

#### 3.3.1. Survival in the Simulated GI Conditions

Following sequential exposure to simulated GI conditions, both strains maintained viable cell counts comparable to their initial levels after 2 min of exposure to SSF (Fig. 4a). After exposure to SGF, DOW-5A maintained a viable cell count of 7.70 ± 0.17 log CFU/mL, whereas DOW-10 decreased to 4.41 ± 0.14 log CFU/mL. At this stage, DOW-5A showed survival rates comparable to that of LGG. The viable cell counts of DOW-5A decreased to 5.12 ± 0.03 log CFU/mL, whereas DOW-10 showed a viable cell count of 5.33 ± 0.21 log CFU/mL. OD₆₀₀ results showed that both isolates generally maintained values comparable to those of LGG throughout sequential exposure to the simulated GI conditions.

**Fig. 4.**
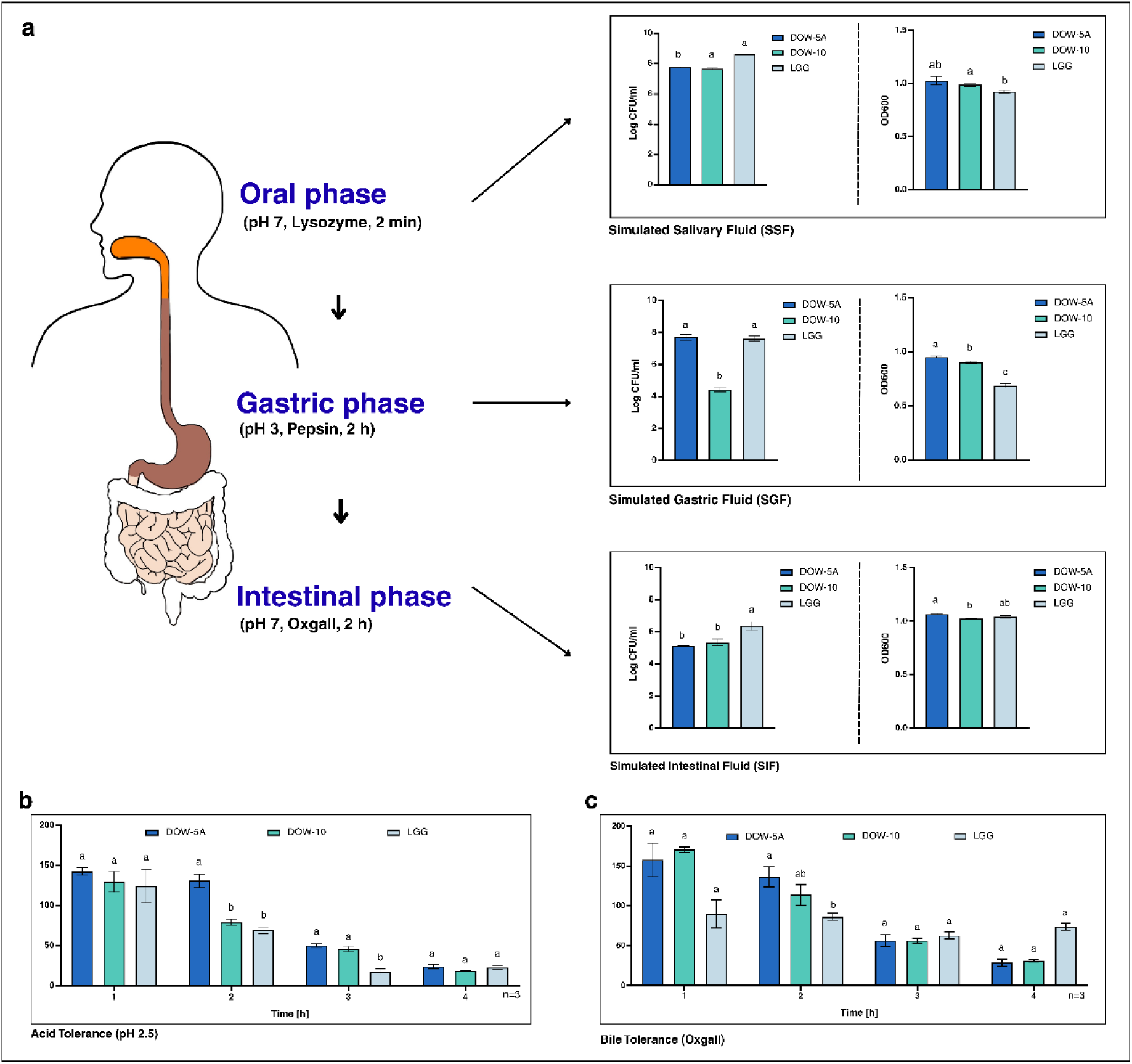
(a) Viable cell counts and OD₆₀₀ were measured during sequential exposure to SSF, SGF, and SIF. Relative growth was assessed under (b) acidic conditions (pH 2.5) and (c) bile conditions (0.3% oxgall) over 4 h. Three independent experiments were performed (n=3). Data were assessed by repeated-measures two-way ANOVA followed by Tukey’s multiple-comparisons test

#### 3.3.2. Acid and Bile Tolerance

The two strains exhibited distinct tolerance patterns over time under acidic (Fig 4b) and bile (Fig 4c) conditions. In pH 2.5 condition, DOW-5A and DOW-10 showed their highest relative growth at 1 h, reaching 142.42 ± 4.73% and 129.63 ± 12.90%, after the values began to decline. DOW-5A maintained high relative growth of 130.91 ± 8.33% at 2 h, exceeding those of the other strains. Under 0.3% oxgall condition, both strains maintained relative growth above 100% during the first 2 h of exposure.

#### 3.3.3. Tolerance to Physiological Stress Conditions

Both strains exhibited concentration-dependent growth under phenol and H₂O₂ stress (Table 2). The relative growth of both strains decreased with increasing phenol and H₂O₂ concentration but remained stable and tended to recover over time. At 6 h, DOW-10 exhibited significantly higher relative growth than LGG under both phenol concentrations. Under 2.0 mM H₂O₂, both isolates exhibited notably reduced relative growth (2–4%) at 6 h, followed by substantially higher values (86–89%) at 12 and 24 h.

**Table 2.** Physiological stress tolerance (phenol and H₂O₂) of DOW-5A and DOW-10

| Physiological stress condition | Concentration | Time (h) | Strain |  |  |
| --- | --- | --- | --- | --- | --- |
|  |  |  | DOW-5A | DOW-10 | LGG |
| Phenol | 0.10% | 6 | 88.48 ± 0.37<sup>b</sup> | 93.82 ± 0.96<sup>a</sup> | 88.24 ± 2.86<sup>b</sup> |
|  |  | 12 | 93.76 ± 1.49<sup>b</sup> | 96.32 ± 7.50<sup>ab</sup> | 96.96 ± 0.42<sup>a</sup> |
|  |  | 24 | 99.67 ± 5.44<sup>a</sup> | 98.41 ± 6.18<sup>a</sup> | 98.08 ± 0.62<sup>a</sup> |
|  | 0.20% | 6 | 58.68 ± 0.82<sup>c</sup> | 73.63 ± 0.77<sup>a</sup> | 69.20 ± 0.76<sup>b</sup> |
|  |  | 12 | 70.06 ± 2.41<sup>b</sup> | 82.88 ± 8.16<sup>a</sup> | 92.98 ± 0.10<sup>a</sup> |
|  |  | 24 | 74.83 ± 9.46<sup>c</sup> | 83.50 ± 7.32<sup>b</sup> | 94.98 ± 1.03<sup>a</sup> |
| H<sub>2</sub>O<sub>2</sub> | 1.0 mM | 6 | 90.21 ± 0.34<sup>c</sup> | 99.14 ± 0.77<sup>a</sup> | 95.34 ± 0.82<sup>b</sup> |
|  |  | 12 | 101.41 ± 0.86<sup>a</sup> | 98.91 ± 0.49<sup>b</sup> | 98.72 ± 0.35<sup>b</sup> |
|  |  | 24 | 97.69 ± 2.53<sup>b</sup> | 94.09 ± 0.18<sup>c</sup> | 98.72 ± 0.20<sup>a</sup> |
|  | 2.0 mM | 6 | 2.75 ± 0.45<sup>b</sup> | 3.87 ± 0.49<sup>b</sup> | 62.56 ± 1.52<sup>a</sup> |
|  |  | 12 | 89.27 ± 0.74<sup>b</sup> | 87.35 ± 0.27<sup>c</sup> | 92.22 ± 0.30<sup>a</sup> |
|  |  | 24 | 89.01 ± 4.36<sup>a</sup> | 86.25 ± 0.34<sup>b</sup> | 96.32 ± 0.43<sup>a</sup> |

Tolerance of DOW-5A and DOW-10 to phenol (0.1–0.2%) and H<sub>2</sub>O<sub>2</sub> (1.0–2.0 mM) conditions was evaluated at 6, 12, and 24 h. The experiments were independently performed three times, with each
sample measured in triplicate in each experiment (n=3). Data were analyzed by repeated-measures two-way ANOVA followed by Tukey's multiple-comparisons test.

### 3.4. Adhesion and Cell Surface Properties

#### 3.4.1. Mucin Adhesion

DOW-5A and DOW-10 showed different adhesion patterns at the two mucin concentration (Fig. 5a). In particular, DOW-5A exhibited the highest adhesion rates of 16.60 ± 4.86% and 15.20 ± 3.32% at mucin concentrations of 0.5% and 1.0%. At 1.0% mucin concentration, DOW-5A exhibited a significantly higher adhesion rate than LGG (P<0.001). These results indicate that both strains were capable of adhering to mucin.

**Fig. 5.**
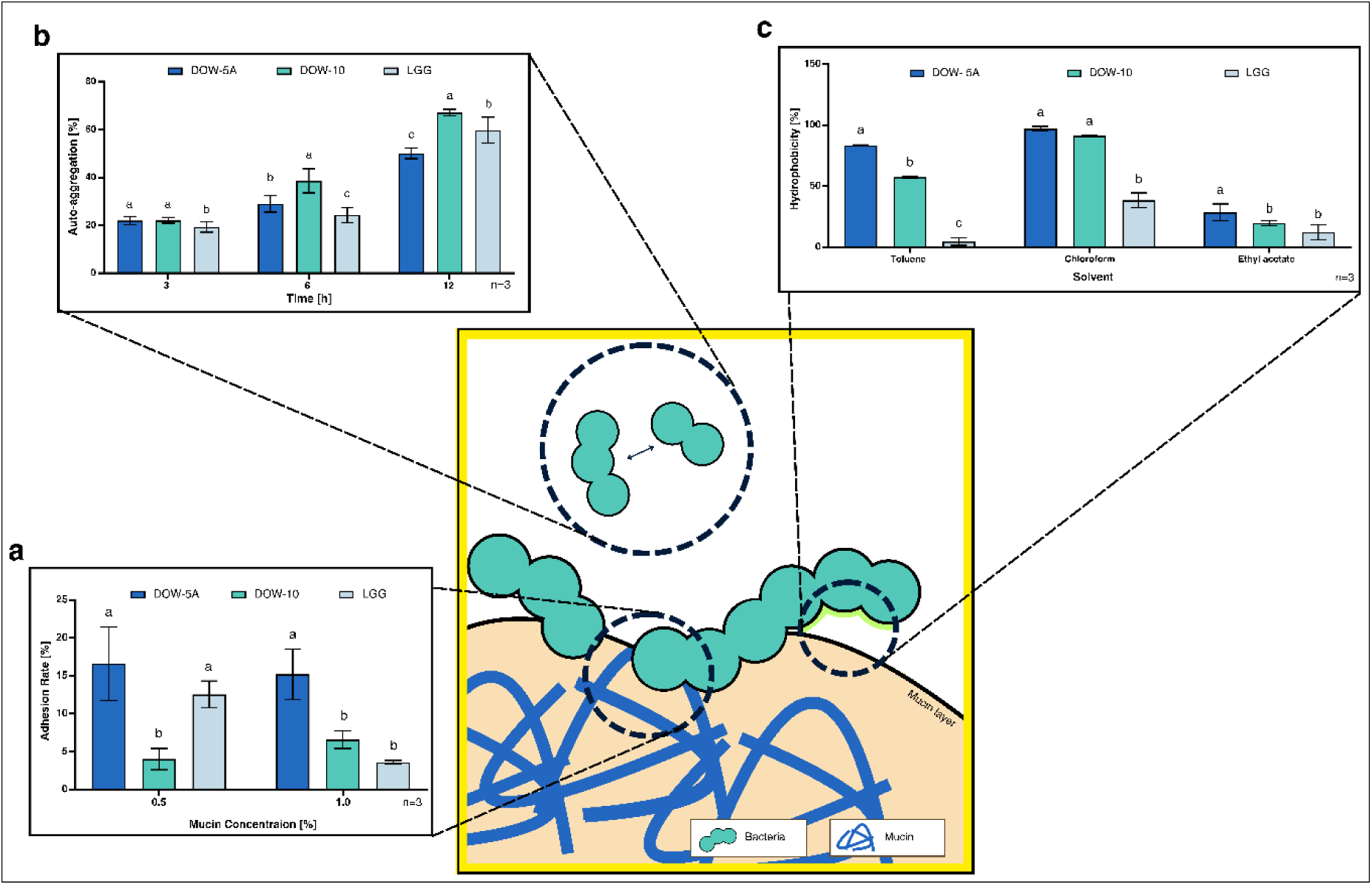
(a) Mucin adhesion at 0.5% and 1.0% (w/v) concentration. (b) Autoaggregation at 3, 6, and 12 h. (c) Cell-surface hydrophobicity toward toluene, chloroform, and ethyl acetate. Three independent experiments were performed for (a) and (c). For (b), three independent experiments were conducted, with each sample measured in triplicate within each experiment (n = 3). Data were analyzed by two-way ANOVA for (a), (c) and repeated-measures two-way ANOVA for (b), followed by Tukey’s multiple-comparisons test

#### 3.4.2. Autoaggregation

Autoaggregation of both strains increased with incubation time, reaching 50% for DOW-5A and 67% for DOW-10 at 12 h (Fig. 5b). DOW-10 exhibited significantly higher autoaggregation than DOW-5A and LGG at both 6 and 12 h (P < 0.01).

#### 3.4.3. Cell Surface Hydrophobicity

Cell-surface characteristics were evaluated by adhesion to different organic solvents (Fig. 5c). DOW-5A exhibited the highest affinity for toluene (83.6 ± 0.4%), chloroform (97.07 ± 1.90%), and ethyl acetate (28.43 ± 6.94%).

### 3.5. Functional Properties

#### 3.5.1. EPS Production

Both strains exhibited EPS-suggestive phenotypes in MRS agar or MRS broth, supplemented with 5% sucrose under both agar and broth culture conditions (Fig. 6a, Supplementary Figure S2). On MRS agar supplemented with sucrose, both strains exhibited a non-mucoid, crystallized colony morphology associated with EPS production.

**Fig. 6.**
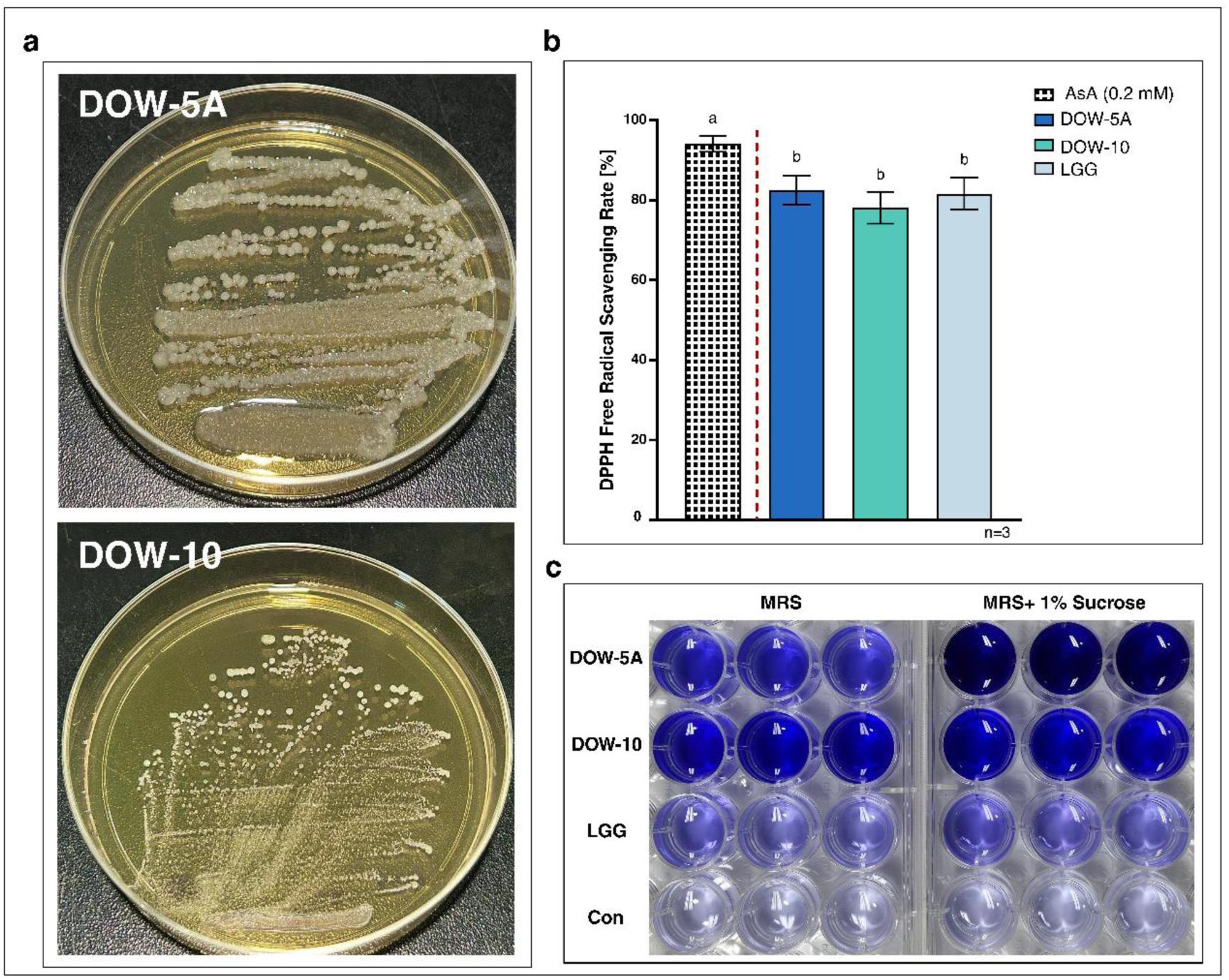
(a) Qualitative assessment of EPS production in MRS medium supplemented with 5% sucrose. (b) DPPH radical scavenging activity with ascorbic acid (0.2 mM) as the positive control. (c) Visual assessment of biofilm formation in MRS medium with or without 1% sucrose following CV staining. The experiments were independently performed three times, with each sample measured in triplicate in each experiment (n=3). Data were assessed using one-way ANOVA followed by Tukey’s multiple comparisons test. AsA: ascorbic acid

#### 3.5.2. DPPH Radical-Scavenging Activity

Both strains exhibited relatively high radical scavenging activity (Fig. 6b). DOW-5A and DOW-10 showed radical scavenging activities of 82% and 78%, comparable to that of LGG (P > 0.05).

#### 3.5.3. Biofilm Formation

Biofilm formation varied among strains in response to sucrose supplementation, with both isolates exhibiting greater biofilm formation than LGG (Table 3). In MRS alone, DOW-10 showed the highest biofilm formation, with a mean corrected OD₅₇₀ of 0.1297. In MRS supplemented with 1% sucrose, DOW-5A exhibited the highest biofilm formation, with a mean corrected OD₅₇₀ of 0.8871. In MRS supplemented with 1% sucrose, DOW-5A exhibited significantly greater biofilm formation than LGG (P < 0.0001). Biofilm formation by DOW-5A reached 1,195% of that in MRS alone, representing the greatest relative increase among the strains following sucrose supplementation. A clear difference was also evident upon visual inspection (Fig. 6c).

**Table 3.** Biofilm formation by DOW-5A and DOW-10

| Strain | Corrected OD<sub>570</sub> |  | Biofilm formation relative to MRS control (%) |
| --- | --- | --- | --- |
|  | MRS | MRS + 1% sucrose |  |
| DOW-5A | 0.0742 ± 0.01<sup>ab</sup> | 0.8871 ± 0.08<sup>a</sup> | 1195.0157 ± 113.79<sup>a</sup> |
| DOW-10 | 0.1297 ± 0.01<sup>a</sup> | 0.1671 ± 0.02<sup>b</sup> | 128.8615 ± 17.08<sup>b</sup> |
| LGG | 0.0191 ± 0.00<sup>b</sup> | 0.0214 ± 0.01<sup>c</sup> | 112.3909 ± 52.60<sup>b</sup> |

Biofilm formation by DOW-5A and DOW-10 was evaluated in MRS broth with or without 1% sucrose using corrected OD<sub>570</sub> values, with relative biofilm formation calculated against MRS broth. The experiments were independently performed three times, with each sample measured in triplicate in each experiment (n=3). Data were assessed by two-way ANOVA for corrected OD<sub>570</sub> and one-way ANOVA for relative biofilm formation, followed by Tukey's test.

#### 3.5.4. Enzyme Activity

Both strains exhibited amylase and protease activities (Supplementary Figure S3). In the amylase assay, DOW-5A and DOW-10 produced clear zones of 18.67 ± 0.58 mm and 11.33 ± 0.58 mm in diameter. In the protease assay, the corresponding clear-zone diameters were 14.33 ± 0.58 mm and 17.00 ± 1.00 mm.

## 4. Discussion

In this study, two wildflower honey-derived *Strep. salivarius* strains exhibited notable responses to GI and physiological stresses, together with a broad range of traits supporting their probiotic potential. Generally, LABs are considered safe, but some species can cause concerns. In our study, both strains were negative for gelatinase and hemolytic activity. Live/dead staining also revealed predominantly live cells, with no distinct increase in dead cells (Fig. 3a), and neither treatment group showed a significant decrease in Caco-2 cell viability in the CCK-8 assay (Fig. 3b). The antibiotic susceptibility profiles of the strains were assessed by measuring their MICs, and differences were observed for vancomycin and erythromycin (Table 1). However, interpretation of the MIC values obtained in this study requires consideration of applicable criteria, and these results alone are insufficient to establish the safety of the strains. Furthermore, MIC testing is recognized for providing a phenotypic assessment that can be influenced by testing conditions and does not directly assess the transfer of resistance genes [45]. Future studies should conduct whole-genome sequencing (WGS) to identify antimicrobial resistance genes and further assessment of their potential for horizontal transfer [46].

For orally administered probiotics, enough viable cells must reach the gut, and survival in the GI transit is an important criterion for evaluating potential probiotic strains [47]. The modified *in vitro* model used in this study assessed this property under sequential exposure conditions. The INFOGEST model is generally used to simulate the digestion of food matrices [35], yet their limitations include an insufficient simulation of the dynamic processes occurring in the actual GI tract and an inability to account for age-related physiological differences [48,49]. Therefore, the intestinal phase in this study contained 0.3% oxgall, intended primarily to strictly evaluate tolerance to acid and bile-related stress rather than to reproduce all components of the standard INFOGEST 2.0 protocol. We also included lysozyme and pepsin as additional stressors. Viable cells of both strains were recovered after sequential exposure, although two strains exhibited distinct survival patterns across the individual phases (Fig. 4). During the gastric phase, DOW-5A maintained viable cell counts comparable to those of LGG. In the separate acid-tolerance assay, its OD₆₀₀ remained relatively stable during the first 2 h at pH 2.5. In contrast, DOW-10 showed a reduction in viable cell counts in the SGF, followed by an increase in the SIF. This increase can reflect the recovery or growth of cells that survived gastric exposure. DOW-5A showed a decrease in viable cell counts during the intestinal phase, which appeared to contrast with its favorable response in the separated assay for oxgall exposure. These results may be attributed to the influence of prior SGF exposure on the subsequent response to intestinal conditions. The separate oxgall assay evaluated an OD₆₀₀-based growth response under a single stress condition, whereas the sequential model quantified viable cells by CFU counting after exposure to oral and gastric stresses. Given these differences in both measurement method and stress exposure, the results of the two assays are not directly comparable. Phenol and oxidative stress are also physiological challenges associated with the GI environment [50,51]. Oxidative stress can occur throughout the gastrointestinal tract, whereas exposure to phenolic compounds is particularly relevant in the colon, highlighting the importance of tolerance to these stresses [52]. Under these conditions tested, the growth of both strains was initially inhibited but increased with prolonged incubation (Table 2). Although this pattern suggests that the inhibitory effect declined over time, OD₆₀₀ measurements alone are insufficient to determine whether this was the result of adaptation or a recovery in growth.

Both strains also exhibited remarkable DPPH radical-scavenging activity comparable to that of LGG, supporting their antioxidant potential (Fig. 6b). The antioxidant capacity of probiotics may contribute to various health-promoting effects, such as the inhibition of lipid peroxidation and the suppression of inflammation by alleviating oxidative stress [53]. Together, the responses to H₂O₂ exposure and the DPPH radical-scavenging activity support complementary aspects of oxidative stress tolerance and antioxidant potential in DOW-5A and DOW-10 [54]. Recent studies report that LAB-derived EPS plays a key role in cell protection and biofilm formation [55]. It has been reported that the EPS produced by *Strep. salivarius* ATCC 9759 during sucrose metabolism via fructosyltransferase (FTF) protected cells from osmotic, acid, and oxidative stress [56], while the water-insoluble glucan and levan produced by *Strep. salivarius* SY511 have been reported to exhibit prebiotic, anti-inflammatory, and antioxidant activities [41]. Given that LAB-derived EPS is well known to contribute to resistance to osmotic stress and biofilm formation [57], this suggests that the EPS produced by the strains may have contributed to their survival in the harsh environment of honey. In our experiments, both strains exhibited characteristic morphological features associated with EPS production (Fig. 6a). Furthermore, DOW-5A exhibited distinct biofilm formation patterns in response to sucrose supplementation (Table 3). The addition of 1% sucrose increased DOW-5A biofilm formation by approximately 12-fold compared with MRS alone. Considering these results together with the EPS-related phenotypes observed in sucrose-containing media, the pronounced sucrose-dependent biofilm formation of DOW-5A could have been associated with EPS production. In contrast, DOW-10 exhibited relatively high biofilm formation even in the absence of sucrose but showed only a limited increase following sucrose supplementation. These findings suggest that biofilm formation and the response to sucrose differed between the two strains. However, because EPS production was evaluated qualitatively based on morphological characteristics in this study, further analysis is required to determine the quantity and composition of the EPS and its direct relationship with biofilm formation [58].

The two probiotic candidates are evaluated not only for their ability to survive GI transit and various physiological stresses, but also for their adhesion to the mucus layer, which is considered an important characteristic related to transient colonization in the gut [59]. In this study, we focused on the direct adhesion to mucin, a major component of the intestinal mucus layer, and evaluated autoaggregation and cell-surface hydrophobicity as adhesion-related surface characteristics. DOW-5A exhibited significantly higher adhesion than LGG and DOW-10 at both mucin concentrations and maintained a similar adhesion rate in higher mucin concentrations (Fig.5a). Both strains showed high adhesion rates at both mucin concentrations, indicating that both isolates were capable of interacting with mucin. Overall, the two strains exhibited differences in adhesion-related characteristics. DOW-5A showed greater affinity for toluene than DOW-10, whereas both strains adhered more strongly to chloroform than LGG (Fig.5c). Adhesion to ethyl acetate was comparatively low, with only DOW-5A showing a significantly higher value than LGG. Together with the pronounced time-dependent autoaggregation observed in DOW-10 (Fig. 5b), these findings suggest that strain-specific differences in cell-surface physicochemical properties may contribute to their distinct adhesion-related phenotypes. In the additional assays for functional characterization, both strains exhibited amylase and protease activities, and TLC analysis qualitatively indicated slight GABA production (Supplementary Figure S4). Tyrosinase inhibitory activity was also observed, but further validation is required because the absence of a positive control limits interpretation of this finding (Supplementary Figure S5). These results indicate that both strains may possess various functional properties beyond the major probiotic characteristics evaluated in this study.

## 5. Conclusion

In conclusion, This study revealed the promising probiotic potential of two Strep. salivarius strains isolated from wildflower honey through a series of assays. Both strains demonstrated diverse functional properties, including tolerance to gastrointestinal conditions and physiological stress, as well as antioxidant activity, biofilm formation, and adhesion-related activities. These findings highlight their potential as probiotic candidates and the strain-specific variability in functional properties. Further genomic characterization and functional validation beyond the present *in vitro* assays are needed to more comprehensively assess their safety and clarify the mechanisms underlying these phenotypes.

## Acknowledgements

The authors thank In-Woo Jung and members of the Department of Life Science, Handong Global University, including Jung-Min Lee, Hongsup Yoon, Bobae Kim, Minseo Hong, and Kyung-Bo Seo, for their valuable support and assistance throughout this study. The authors also thank Hanbyoul Jung for English-language editing.

## Statements and Declarations

### Compliance with Ethical Standards

#### Ethical Approval

Ethical approval was not required for this study, as only an established human cell line was used and no human participants or newly collected human biological materials were involved.

#### Data Availability

The data supporting the findings of this study are available from the corresponding author upon reasonable request. The 16S rRNA gene sequences of strains DOW-5A and DOW-10 have been deposited in the NCBI GenBank database under accession numbers PZ626003 and PZ626004 and will be publicly available upon publication.

#### Author contributions

Wonhui Lee: Investigation, Conceptualization, Formal analysis, Project administration, Funding acquisition, Validation, Writing – original draft, Writing – review & editing. Yeon-Soo Oh: Investigation, Formal analysis, Data curation, Validation, Writing – original draft, Writing – review & editing. Chan-Soo Ock: Funding acquisition, Validation, Writing – review & editing. Ye-Eun Oh: Validation, Writing – review & editing. Ha-young Kim: Investigation. Hyung-Ki Do: Conceptualization, Project administration, Funding acquisition, Validation, Supervision, Writing – review & editing. Chul-Won Hwang: Resources, Writing – review & editing. All authors read and approved the final manuscript. W.L. and Y.-S.O. contributed equally to this work and share first authorship.

## Supplementary Figure

**Supplementary Figure S1.**
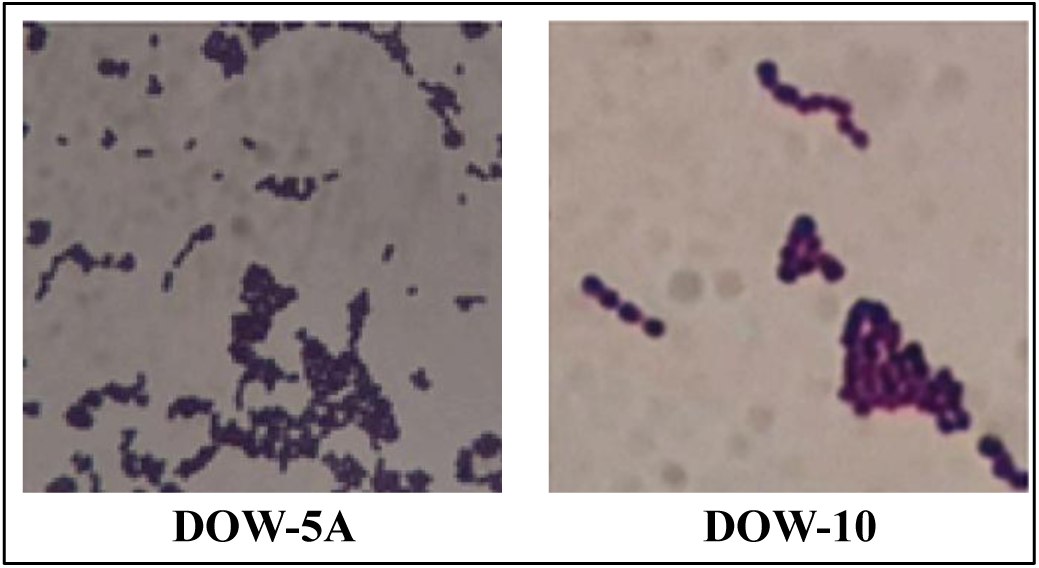
Gram-staining morphology of DOW-5A and DOW-10.

**Supplementary Figure S2.**
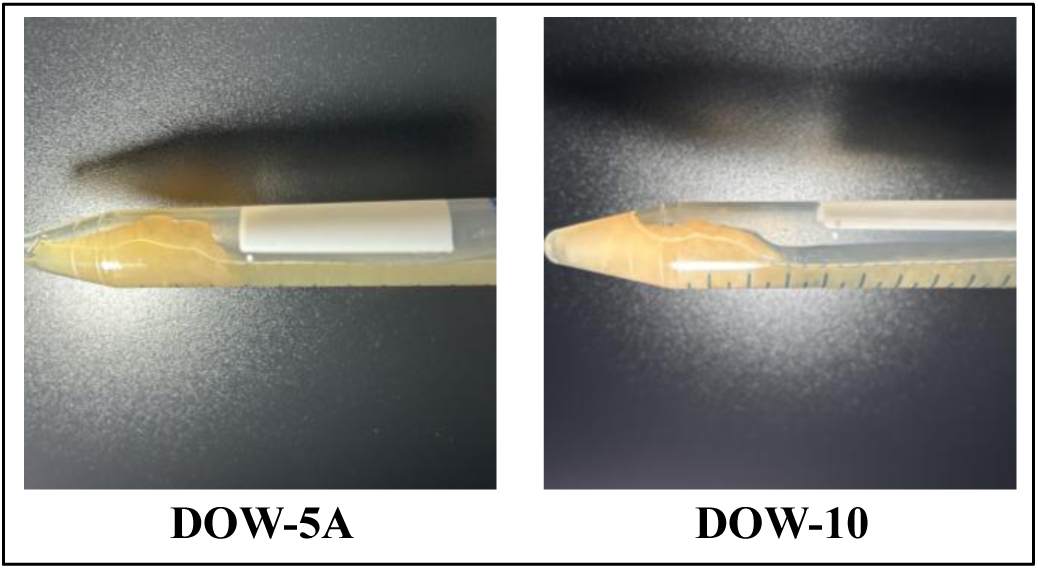
EPS-related phenotype of DOW-5A and DOW-10 in 5% sucrose supplemented MRS broth.

**Supplementary Figure S3.**
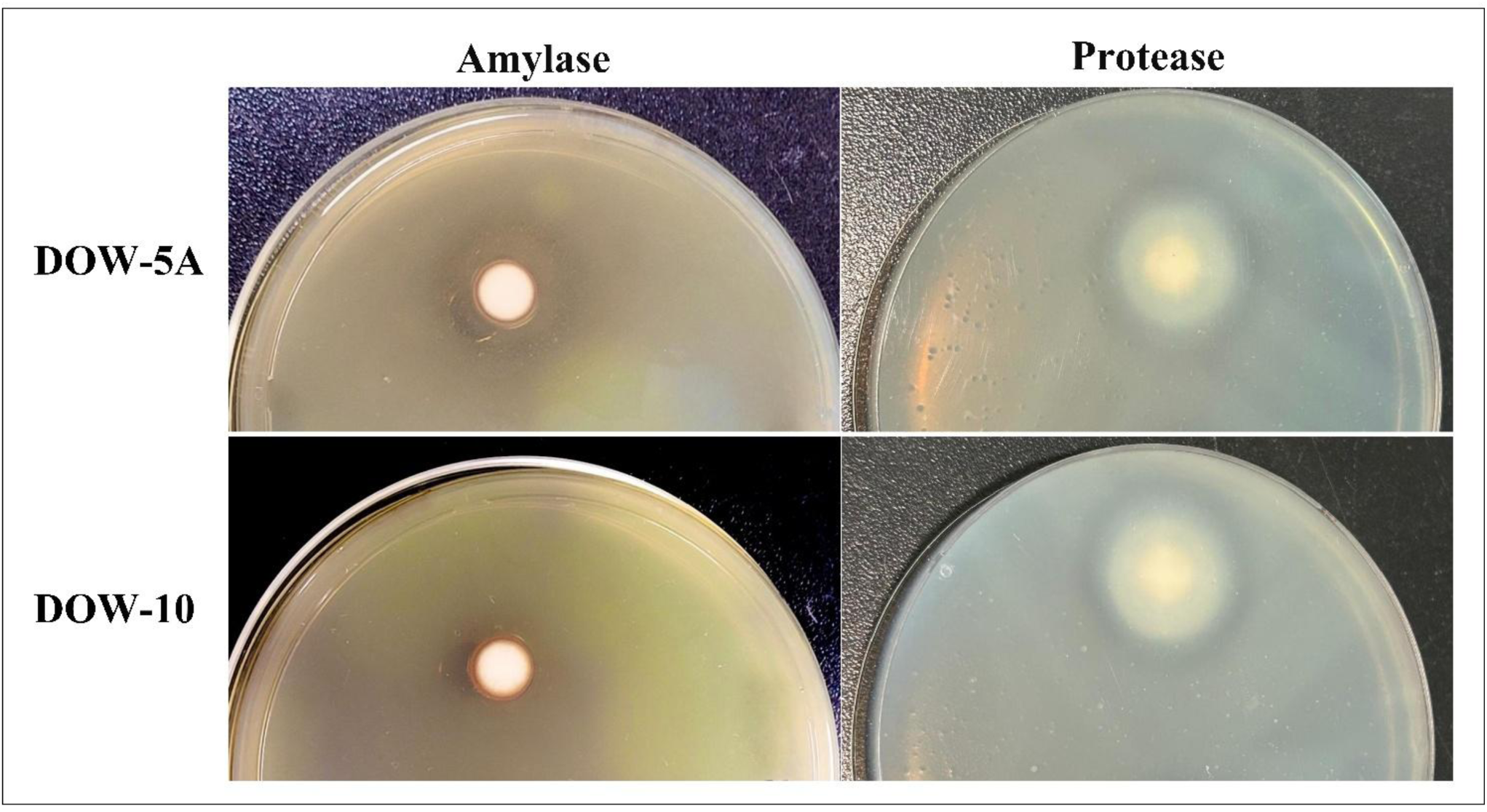
Amylase and protease activities of DOW-5A and DOW-10.

**Supplementary Figure S4.**
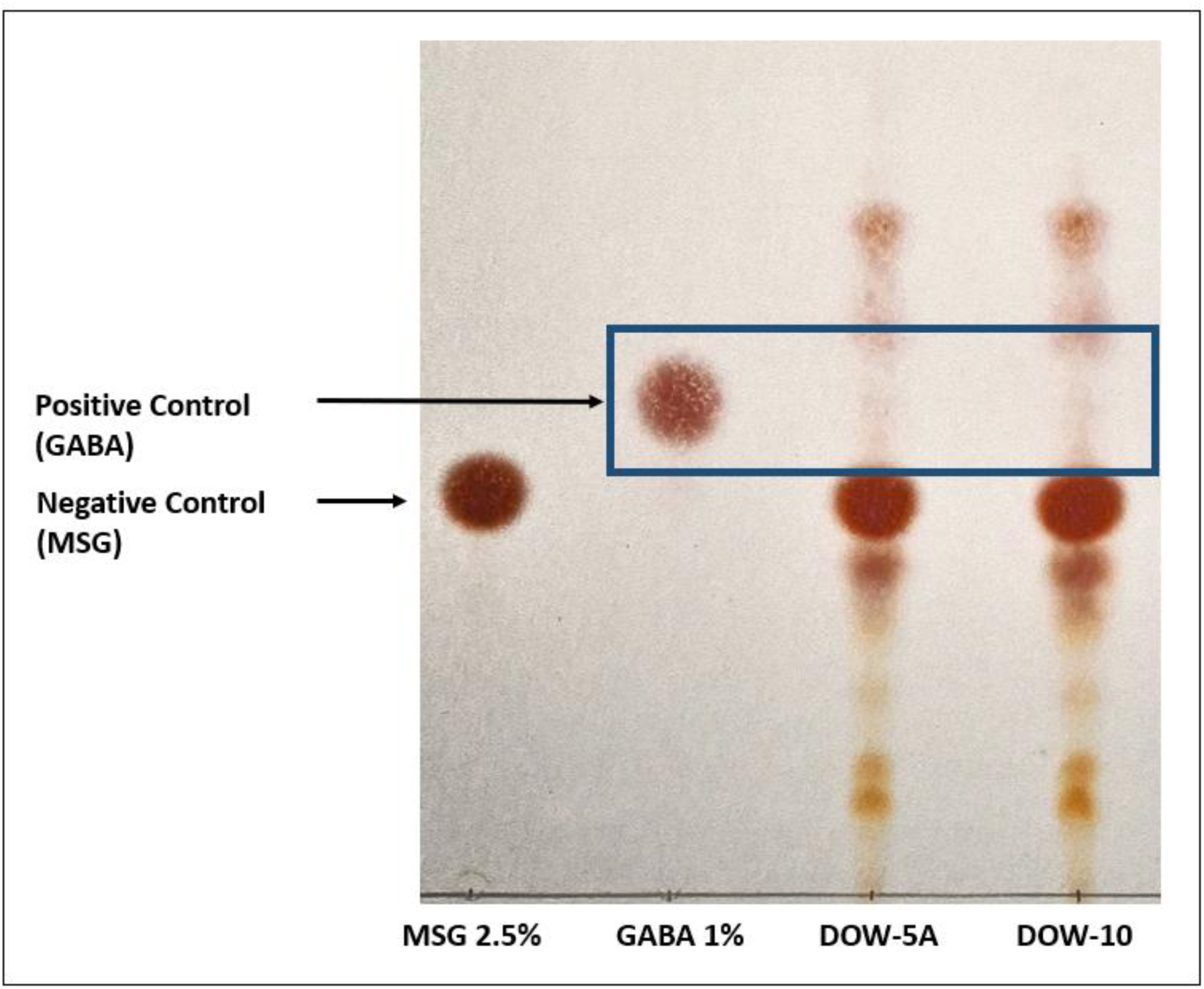
Qualitative analysis of GABA production by DOW-5A and DOW-10 using thin-layer chromatography (TLC). 1% GABA standard was used as a reference for the identification of corresponding spots in the bacterial samples.

**Supplementary Figure S5.**
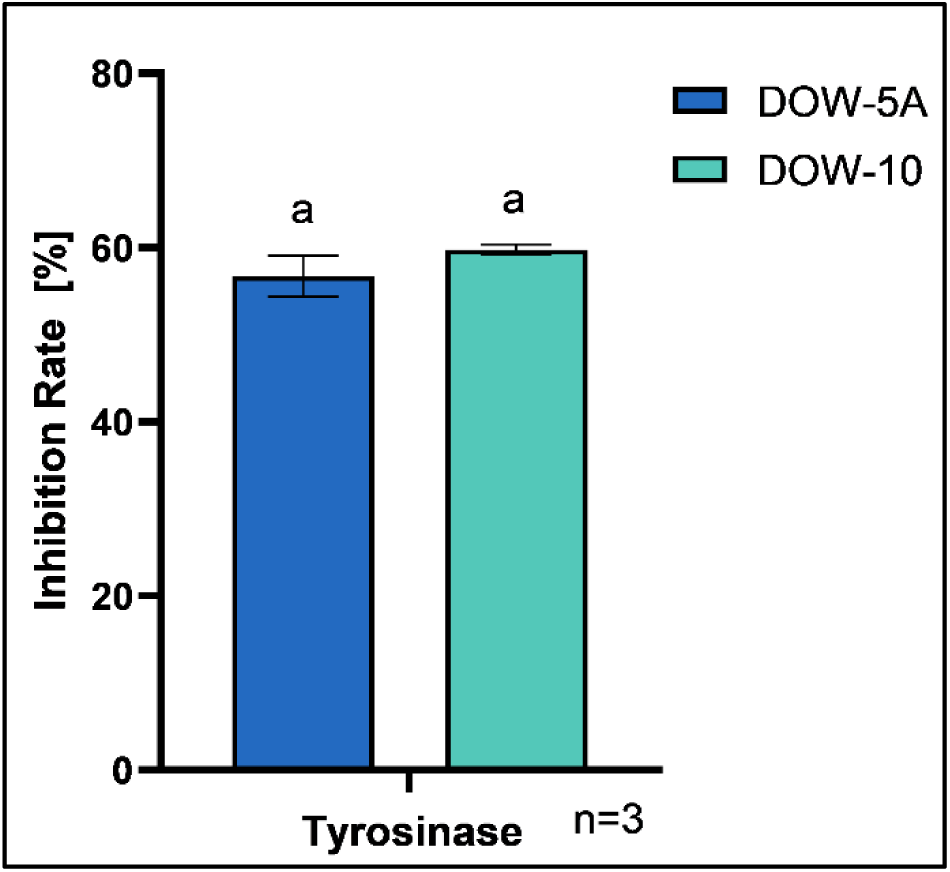
The tyrosinase inhibitory activity of DOW-5A and DOW-10 was evaluated using mushroom tyrosinase with L-tyrosine as the substrate. Statistical significance was analyzed using an unpaired two-tailed Student’s t-test to compare DOW-5A and DOW-10.

## Supplementary Method

### GABA production

GABA production was conducted with modifications based on the protocol [60]. Cultures grown at 37 °C for 24 h were inoculated at 1% (v/v) into MRS broth supplemented with 2.5% (w/v) sodium L-glutamae (Daejung Chemicals and Metals) and incubated at 37 °C for 4 days. Culture supernatants (1 μL) were spotted onto silica gel TLC plates (Merck Millipore, Darmstadt, Germany) and developed for 2 h with *n*-butanol (Daejung), acetic acid (Daejung), DW (3:2:1, v/v/v). 1% (w/v) GABA solution was used as the standard. After development, the plates were dried, sprayed with 0.2% (w/v) ninhydrin (Sigma-Aldrich) in ethanol, and heated at 80 °C. GABA production was identified by comparison with the GABA standard (Sigma-Aldrich, St. Louis, MO, USA).

### Tyrosinase inhibition

Cultures grown at 37 °C for 24 h were centrifuged at 8,000 × g for 2 min at 4 °C. 300 μL of supernatant was mixed with 225 μL of 0.4 M HEPES buffer (pH 6.8), 150 μL of mushroom tyrosinase (150 U/mL), and 225 μL of 2.5 mM L-tyrosine to a final volume of 900 μL. In the control, the supernatant was replaced with HEPES buffer. After incubation at 30 °C for 15 min, absorbance was measured at 475 nm. Tyrosinase inhibition was calculated as follows [61]:

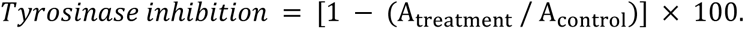

A_treatment_ and A_control_ represent reactions with and without culture supernatant.

